# MOLECULAR INSIGHTS INTO PESTICIDE TOLERANCE: PROTEIN INDUCTION AND DNA DAMAGE IN ORGANOPHOSPHATE-DEGRADING BACTERIA

**DOI:** 10.64898/2026.04.25.720663

**Authors:** Neethu Asokan

## Abstract

One of the effects of the intensified agricultural activities involves environmental pollution by pesticides, which are bound to get into the soil and ultimately into the water sources through leaching. The recurrent exposure of soil microbiota to these poisonous substances facilitates the process of adaptive resistance and catabolic functions. In the current research, bacterial cultures taken in Karuppur and Salem pesticide-contaminated agricultural soils were filtered on their capability to decompose organophosphate pesticides. Two strong isolates, which were referred to as *Bacillus sp*. and *Micrococcus sp*. had a great level of tolerance and degradation capacity. Significant biomolecular changes in these isolates were observed after long-term exposure (three months) to organophosphate pesticides. A protein estimation showed a strong rise in the overall total protein content indicating the activation of stress-related and degradative enzymes. Genomic DNA damage was identified by DNA ladder assay, which is a genotoxic stress caused by pesticides. Thus, plasmid profiling also revealed a rise of copy number and change of the size of plasmids, implying potential adaption through plasmids and greater degradation potential. This evidence indicates that long-term exposure to pesticides leads to microbial adaptation in terms of physiological and genetic changes to allow survival in adverse environments. The isolates identified have great potential to be used in bioremediation strategies that will be used in detoxifying the soils that have been contaminated with organophosphate.

## 1. Introduction

In India, agriculture is one of the major sources of livelihood and almost ~70 percent of the population depends on it either directly or indirectly. To sustain the growing food demand, application of chemical pesticides has become a part and parcel of contemporary agricultural activities. Agrochemicals such as herbicides, insecticides and others have greatly enhanced the quality and quantity of crops while still making them economically viable. However, their indiscriminate and prolonged application has led to serious environmental and health concerns. These include organophosphorus pesticides (OPs) which are highly used because they are very effective against a broad variety of insect pests. Some of the common OPs used including Chlorpyrifos, Parathion, Monocrotophos and Dichlorvos are used extensively in agricultural systems. The toxicity of these compounds is mainly by inhibiting the activity of acetylcholinesterase thus interfering with the transmission of nerve impulses by the target and non-target organisms (Tudi et al., 2023; Cheng et al., 2025). As a result, organophosphate poisoning has become one of the most significant worldwide health challenges with 3 million cases reported annually (Chukwu et al., 2025).

The large application of pesticides has led to contamination of soil and groundwater systems by leaching, runoff, and settling. This environmental conservation does not only interfere with the soil fertility but also with the microbial diversity and ecology (Yasemi et al., 2025). Moreover, residues of pesticides may cause long-term health consequences, such as neurological, endocrine, reproductive, and immunological side effects, and carcinogenic effects to human beings (Okolonkwo et al., 2022). Chen et al., (2024) have also reported mechanisms of neurotoxicity caused by organophosphate pesticides in humans.

Pesticides may also be utilized as food by the microbes even though they are toxic. Microorganisms of soil, especially bacteria, are important in the biodegradation and detoxification of the xenobiotic compounds. Organophosphate pesticides have been reported to be degraded by several genera of bacteria, such as Pseudomonas, Bacillus, and Arthrobacter, which use them as either a carbon or phosphorus source (Anjum et al., 2022). This microbial adaptation is also commonly induced by the long-term exposure that favours resistant populations that can metabolize toxic compounds (Dhakal et al., 2025).

Moreover, ongoing exposure to pesticides has been demonstrated to cause physiological and genetic alterations of microbial communities such as increased enzyme synthesis, stress response systems and transfer of plasmid genes (Yasemi et al., 2025). These adaptations can also potentially cause cross resistance effects, such as resistance to antibiotics increasing as seen in bacteria that are exposed to herbicides (Zhao et al., 2026). Due to the growing environmental impact of pesticides contamination, there is increased interest in studying indigenous microbial populations and their potential use in the context of bioremediation. The current research is based on the isolation, screening and characterization of organophosphate-degrading bacteria of pesticide-treated agricultural soils. In addition, the paper measures their tolerance to different pesticides in different concentrations and examines the effects of long-term exposure on the expression of proteins and DNA integrity and plasmid profiles.

## 2. Materials and Methods

### 2.1. Sample Collection

The agricultural fields around Periyar University, Salem, Tamil Nadu, India were sampled to obtain soil samples. Sterile polythene bags were used to collect samples in a sterile environment and take them to the lab where they were further analyzed.

### 2.2. Isolation of Bacteria

A 100 mL of sterile distilled water was added to a 1 gram of each soil sample to achieve an initial dilution (10 −2). Serial dilutions were done up to 10 7 by adding 1 mL of the preceding dilution to 9 mL of sterile diluent. Aliquots (0.1 mL) of 10 −4, 10 −5 and 10 −6 were placed on sterile nutrient agar plates. The incubation was carried out at 37 o C. Separate colonies were picked and kept in nutrient agar slants to be used in further investigations.

### 2.3. Screening of organophosphate-degrading bacteria (Elçin, 2025)

The bacterial isolates were selected based on their capability to degrade the organophosphate pesticides with the help of Mineral Salt Medium (MSM) nutritionally supplemented with 0.1% organophosphate pesticides, i.e., Monocrotophos and Dichlorvos. Cultures were cultivated at 37C in 48 h with occasional shaking. The cultures were then incubated after which the cultures were transferred to nutrient agar plates that had been supplemented with 0.1% pesticides. Isolates that exhibited growth were chosen to be further characterized.

### 2.4. Characterization of Bacterial Isolates

The morphological and biochemical characterization of the isolates selected were done using standard procedures in microbiology (Barahmbhatt, 2014). The tests were Gram staining, motility test, catalase test, oxidase test, endospore staining, citrate utilization, urease activity, production of hydrogen sulfide, and utilization of carbohydrates (glucose and lactose).

#### 2.4.1 Gram Staining

A 24 h culture on a clean glass slide was heat-fixed and stained in turn with crystal violet (1 min), Grams iodine (1 min), decolorized with alcohol and stained with safranin (30 45 s). The slides were viewed using oil immersion microscopy.

#### 2.4.2 Motility Test

The hanging drop technique was used to determine motility. An albeit drop of 6-8 h of culture was put on a coverslip and inverted on a concavity slide and observed using a 40x objective.

#### 2.2.3. Catalase Test

One drop of 3-percent hydrogen peroxide was added to a loopful of bacterial culture. The formulation of a bubble was an indication of a positive result.

#### 2.4.4 Oxidase Test

A bacterial culture was put on an oxidase disc, and the appearance of the color change depicted that the bacterium was oxidase positive.

#### 2.4.5 Endospore Staining

Heat-fixed smears were stained with 5% malachite green with 5 min steaming conditions. Washing of slides was followed by counterstaining with safranin (30-60 s) and examining using oil immersion microscopy.

### 2.5 Organophosphate Pesticide Tolerance of Bacterial Isolates

Isolates were selected and tested to determine their tolerance, with different concentrations (0.1, 0.5, 1, 5, 10, 25) of the organophosphate pesticides (monocrotophos and/or dichlorvos). The cultures were incubated during 48 h and the growth of bacteria was spectrophotometrically measured at 620 nm. The most tolerant isolates were picked to continue with further research.

### 2.6 Long-Term Exposure of Bacteria to Pesticides

The bacterial isolates were first cultivated in MSM during the period of 24 hours at 37C, and then transferred into LuriaBertani (LB) medium that contained 0.1% pesticides. Adaptive responses were studied by keeping the cultures three months and adding pesticides periodically at weekly intervals.

### 2.7 Impact of Bacterial Biomolecules on Pesticide Exposure. The total protein was estimated using the lowry method

The Lowry method was used to estimate the total protein content.

Reagents Preparation:

2% Na □ CO□ in 0.1 N NaOH

1% sodium potassium tartrate

0.5% CuSO□·5H□O

Reagent I: Na 2 CO 3, tartrate, and CuSO 4 mixture.

Reagent II: Folin–Ciocalteu reagent (1:1 with distilled water) dilution.

Bovine Serum Albumin (BSA) standard (1 mg/mL).

#### Procedure

To develop a standard curve, standard solutions were prepared with the BSA and the absorbance at 660 nm was measured. In the case of test samples, bacterial cultures which were subjected to pesticides were centrifuged and the supernatant was subjected to reagents I and II, after which, incubation was carried out. The absorbance at 660 nm was measured with a UV Vis spectrophotometer and the concentration of the protein was determined using the standard curve.

### 2.7. The DNA Ladder Assay (Genomic DNA Isolation)

The genomic DNA was isolated using phenol-chloroform technique. SDS was used to lysed bacterial cultures, and phenol-chloroform was used to extract the bacteria and sodium acetate-isopropanol was used to precipitate the bacteria. The DNA pellet was then rinsed with 75 percent ethanol, dried in the air and then again in TE buffer. Electrophoresis on 0.4% agarose gel confirmed the DNA integrity.

#### 2.7.1 Plasmid Isolation

Isolation of plasmid DNA was done by alkaline lysis method. The cells of bacteria were collected, resuspended in solution I, lysed in solution II, and neutralized in solution III. Ethanol was used to precipitate plasmid DNA in supernatant after centrifugation, and washed with 70% ethanol, dried, and dissolved in TE buffer. The plasmid DNA was left alone and kept under −20 C to analyze it.

## 3. Results and Discussion

### 3.1 Isolation of Bacterial Strains from Pesticide-Exposed Soil

Fifty morphologically different bacterial colonies were cultivated out of the agricultural soils that had been subjected to pesticides in 3-5 months. The difference in colony-forming units (CFU) of the samples is an indication that long-term exposure to pesticides influences microbial population dynamics. It is noteworthy that the number of CFU was relatively higher in samples that were longer exposed (45 months), with a maximum of 108 CFU/mL, which suggested the potential enrichment of microbial communities that were resistant to pesticides (Figure 1).

**Figure 1:**
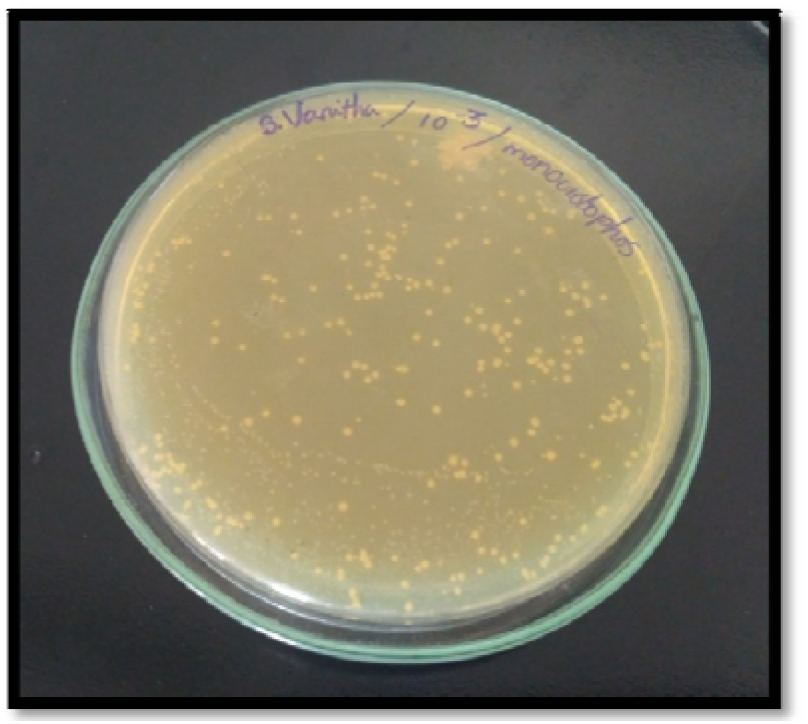
Isolation of potential bacteria.

This finding is like the earlier reports that documented that the unending exposure to pesticides has a selective effect, promoting the increase in the number of resistant or degrading microbial populations (Majumder et al., 2021). This adaptive enrichment is a major ecological behaviour to increase biodegradation capacity of polluted soils. Prashar et al., (2016) have also reported similar reports on the negative impact of pesticides on microbial communities in soil.

### 3.2 Evaluation of the Organophosphate-Degrading Bacteria

Out of the 50 isolates, only 11 of them showed growth in nutrient agar supplemented with 0.1% organophosphate pesticides, which indicates that they can break down organophosphates. The difference in the degree of growth (++ to +++) indicates that there are variations in the metabolic ability of the isolates (Table 4; Figure 6-13).

The capacity of these isolates to grow in pesticide-contaminated media suggests that there is a mechanism of utilization or tolerance, e.g., enzymatic hydrolysis of organophosphate compounds. The breakdown of organophosphates by microorganisms is in most cases catalyzed by enzymes such as organophosphorus hydrolases and esterases, which transform toxic substances into less toxic products (Ghanem et al., 2007). Similar studies were conducted by Eissa et al., (2014) reporting the biodegradation of Chlorpyrifos by microbial communities in soil.

### 3.3 Bacterial Isolates Tolerance to Higher Pesticide Concentrations

The 11 picked isolates were further tested on tolerance to increasing concentration (0.1% - 25%) of organophosphate pesticides such as Monocrotophos and Dichlorvos.

The isolates had a high tolerance with isolate V1, V3 and V5 tolerating up to 5% monocrotophos and V4 and V5 tolerating up to 10% dichlorvos. The displayed reduction of the optical density at the higher concentrations (25%) indicate inhibitory effects because of stronger toxicity (Table 1,2).

**Table 1:** Tolerance of bacterial isolates against different concentration of Monocrotophos (%)

| Sl No | Pesticide<br>(Monochrotophos)<br>% | OPTICAL DENSITY AT 620 NM |  |  |  |  |
| --- | --- | --- | --- | --- | --- | --- |
|  |  | V<sub>1</sub> | V<sub>2</sub> | V<sub>3</sub> | V<sub>4</sub> | V<sub>5</sub> |
| 1 | 0.10% | 0.104 | 0.168 | 0.117 | 0.158 | 0.209 |
| 2 | 0.50% | 0.255 | 0.325 | 0.311 | 0.284 | 0.24 |
| 3 | 1% | 0.816 | 1.178 | 1.467 | 1.167 | 1.363 |
| 4 | 5% | 1.585 | 1.09 | 1.538 | 0.976 | 1.475 |
| 5 | 10% | 0.428 | 1.056 | 1.386 | 0.65 | 1.339 |
| 6 | 25% | 0.119 | 0.609 | 0.439 | 0.309 | 0.591 |

**Table 2.** Tolerance of bacterial isolates against different concentration of Dichlorvos(%)

| Sl.No | Pesticide<br>(Dichlorvos) % | OPTICAL DENSITY AT 620 NM |  |  |  |  |
| --- | --- | --- | --- | --- | --- | --- |
|  |  | V 1 | V 2 | V 3 | V4 | V5 |
| 1 | 0.10% | 0.106 | 0.188 | 0.238 | 0.237 | 0.229 |
| 2 | 0.50% | 0.192 | 0.109 | 0.248 | 0.291 | 0.188 |
| 3 | 1% | 0.445 | 0.81 | 1.193 | 2.103 | 1.607 |
| 4 | 5% | 1.697 | 0.128 | 2.246 | 2.377 | 0.494 |
| 5 | 10% | 2.015 | 2.026 | 2.03 | 2.042 | 2.399 |
| 6 | 25% | 0.13 | 0.984 | 0.267 | 0.686 | 2.018 |

These results are consistent with the previous reports that microbial tolerance to pesticides reduces past the threshold levels due to enzyme inhibition and toxicity of the cells (Hussain et al., 2009; Hussain et al., 2021). Nevertheless, some isolates (especially V5) can withstand higher concentrations, which implies that they may have some efficient detoxification strategies or adaptive stress responses.

### 3.4 Long-term Exposure and Adaptive Stability of Isolates

Pesticides were applied on the selected isolates that were exposed to prolonged exposure (three months). The findings showed a continuous increase in both MSM and LB media with 0.1 percent pesticides, and hence adaptive stability (Figure 15,16).

The prolonged exposure would result in physiological and genetic changes, such as degradation pathways and stress responses systems upregulation. Adaptive patterns have been observed in pesticide-polluted ecosystems where microbial communities have developed increased degradation efficiency (Cycoń et al., 2009). Cycoń et al., (2019) have also reported the degradation and impact of antibiotics on microbial community, their activities and adaptations.

### 3.5 Pesticide Exposure Effect on Protein Expression

The Lowry protein estimation showed that there was an increment in total protein contents in the pesticide exposed isolates as compared to controls. To illustrate, isolate V1 was improved by 0.28 mg/mL (control) to 0.46 mg/mL (treated), and isolates V4 and V5 had higher levels of proteins after being exposed to dichlorvos (Table 3,4).

**Table 3.** Protein concentration of V1 after exposure to Monocrotophos.

| S.NO | OPTICAL<br>DENSITY AT 660<br>NM (V<sub>1</sub>) | CONCENTRATION OF<br>PROTIEN IN mg/ml |
| --- | --- | --- |
| CONTROL |  |  |
| 1 | 0.729 | 0.28 |
| TEST SAMPLE |  |  |
| 1 | 0.931 | 0.46 |

**Table 4.** Protein concentration of V4 and V5 after exposure to Dichlorovos.

| S.NO | OPTICAL DENSITY<br>AT 660 NM | CONCENTRATION OF<br>PROTIEN IN MG |
| --- | --- | --- |
|  | CONTROL |  |
| 1 | 0.832 (V<sub>4</sub>) | 0.33 |
| 2 | 0.605 (V<sub>5</sub>) | 0.24 |
|  | TEST SAMPLE |  |
| 1 | 0.668 (V<sub>4</sub>) | 0.66 |
| 2 | 0.519 (V<sub>5</sub>) | 0.5 |

**Table 5.** Identification study of potent isolate.

| Sl No | Identification test | V1 | V4 |
| --- | --- | --- | --- |
| 1. | Gram staining | Gram positive rod | Gram positive cocci |
| 2. | Motility | Motile | Non motile |
| 3. | Endospore | Positive | Negative |
| 4. | Catalase | Positive | Positive |
| 5. | Oxidase | Negative | Positive |
| 6. | Indole | Negative | Negative |
| 7. | Citrate | Positive | Positive |
| 8. | MR | Negative | - |
| 9. | VP | Positive | - |
| 10. | Starch hydrolysis | Positive | Negative |
| 11. | Selective media |  |  |
|  | a. Mannitol salt agar | Negative | Positive -light yellowish colony |
|  | b. Starch agar | Positive | Negative |

The protein content increase can be explained by the fact that the enzymes of stress-induction and degradation that are related to the metabolism of pesticides are induced. The increased protein synthesis during the stressful conditions is a well-documented stress-response in microbes, which is commonly linked to the synthesis of enzymes to neutralize toxins and protective proteins (Wei et al., 2025).

These findings indicate that isolates V1 and V4 are more metabolically active and have a greater degradation capacity and can be used as a good bioremediation candidate.

### 3.6 Analysis of DNA Damage (DNA Ladder Assay)

Smeared DNA bands were observed in the isolates V1, V4 and V5 after exposure to pesticides, which showed that the DNA was damaged. The DNA fragmentation is a typical result of the oxidative stress caused by toxic substances like organophosphates. The organophosphate pesticides have been reported to produce reactive oxygen species (ROS) that can lead to strand breaks and damage DNA (Altuntas et al., 2003). The smearing of the DNA indicates that even though the isolates can withstand the exposure to pesticides, they are still subjected to genotoxic stress (Figure 17 a,b).

Such a two-fold reaction of tolerance and DNA damage indicates the trade-off between survival and cellular stress in pollutant environments.

### 3.7 Potent Bacterial Isolates

According to morphological and biochemical characterization, isolate V1 was determined to be Bacillus subtilis and isolate V4 as Micrococcus luteus (Table 4; Figure 2-5).

**Figure 2:**
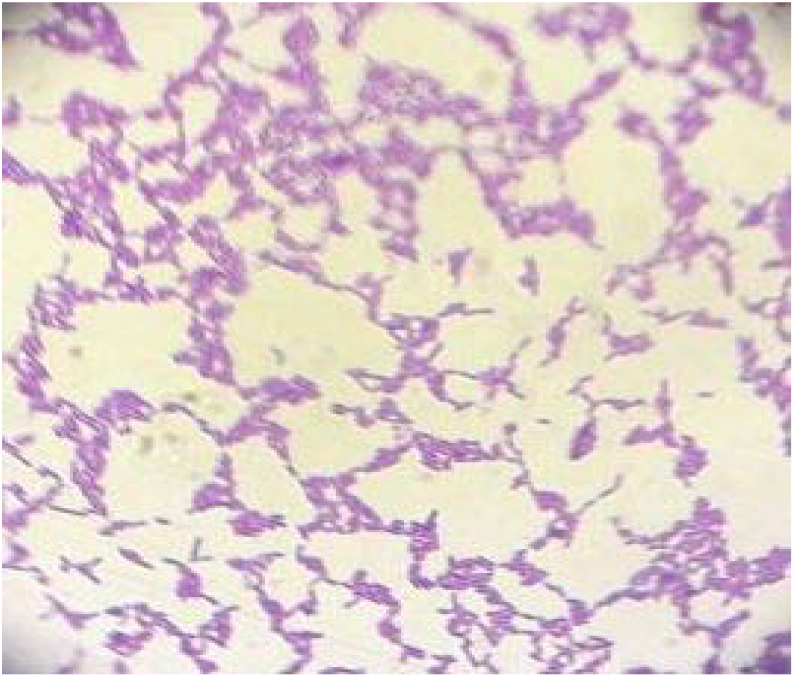
Gram positive rod (V_1_) under 40X microscopic view.

**Figure 3:**
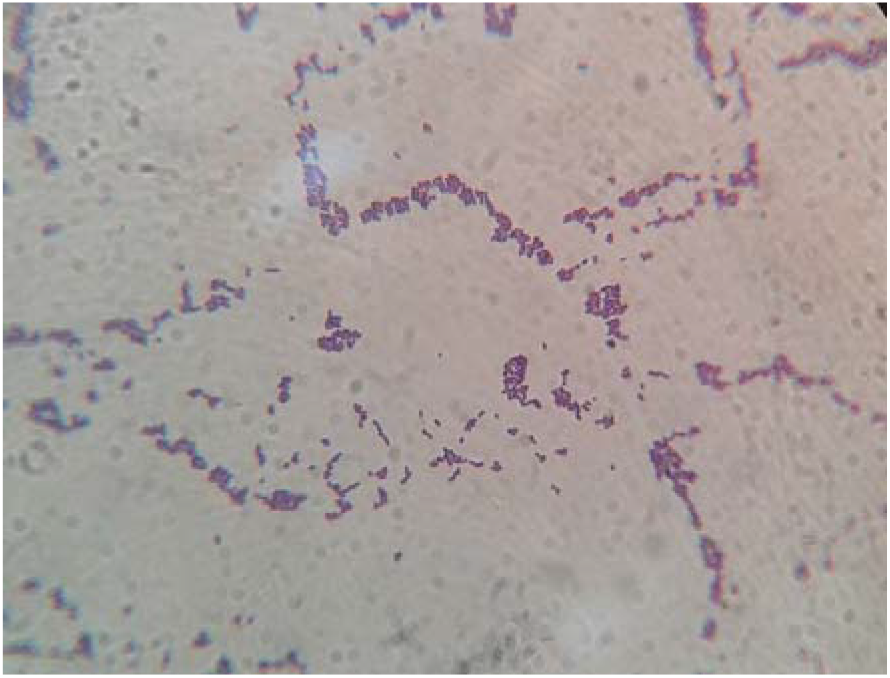
Gram positive cocci.

**Figure 4:**
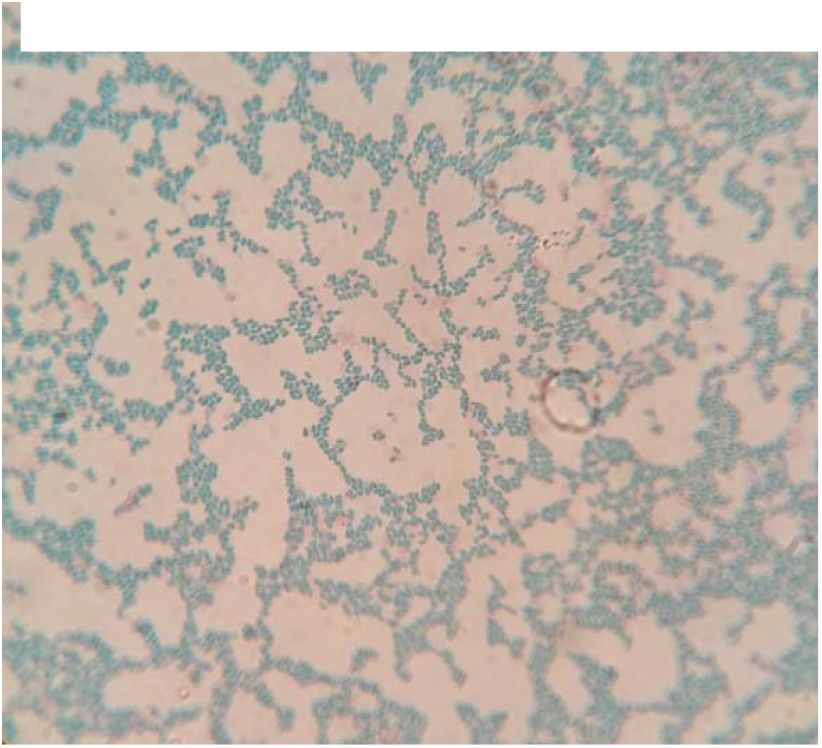
Spores under 100x microscopic view (v1)

**Figure 5:**
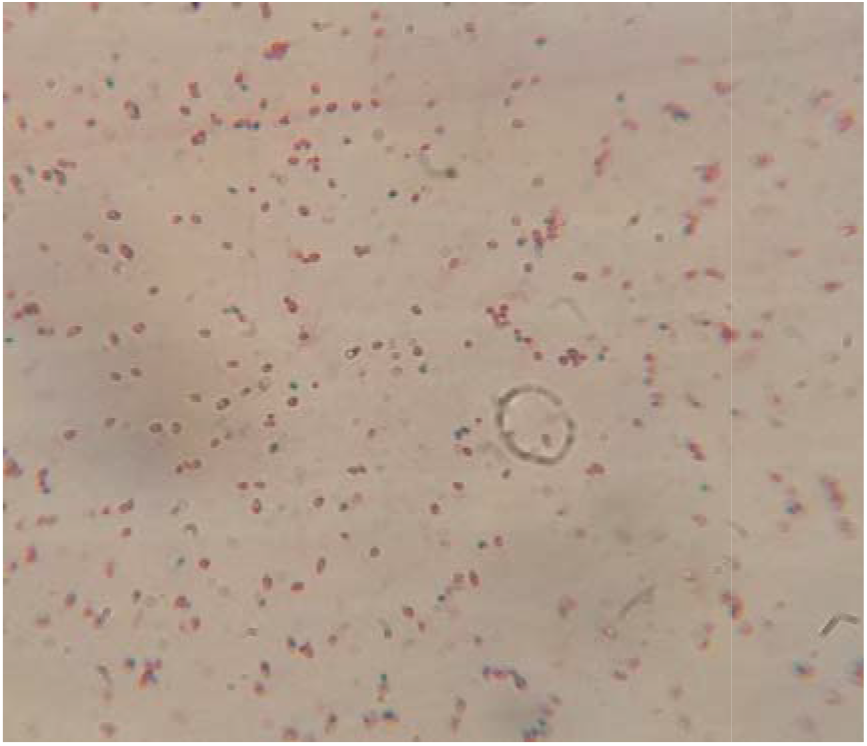
Vegetative cells without spores under 100x microscopic view.

Both species are well documented in terms of environmental resilience and biodegradation. Bacillus subtilis can produce numerous enzymes in the xenobiotic degradation process whereas Micrococcus luteus has been reported to survive in harsh and contaminated environments with effective detoxification systems (Das and Chandran, 2011).

## 4. Conclusion

The research shows that soils that are exposed to pesticides contain a wide variety of bacterial communities capable of high biodegradation. The gradual increase in tolerant strains and improved protein expression, as well as adaptation via plasmid all point to the evolution of efficient microbial systems that can break down organophosphates. These results justify the possibility of using the native bacterial isolates in the sustainable bioremediation processes of pesticides-contaminated environments. The present study gives an insight on the pesticide degrading bacteria and the major changes evolved in the bacteria because of long term pesticide exposure. Soil microbiota acquires several changes in their biomolecule level that enable them to sustain and degrade these toxic compounds. It can be said that the bacteria assimilate and utilize the pesticide thus minimizing the toxic effect. The plasmid changes are one major significant factor that enables the bacteria to either get altered in such a way to overcome the toxicity of the pesticide. These bacteria with several degrading proteins, genes, DNA and plasmid changes could be a better tool for bioremediation of the pesticides. The findings of this study highlight the prospects of these bacteria along with gives an insight on the impact of these pesticides on the bacterial biomolecules.

## ACKNOWLEDGEMENT

Authors acknowledges the Vice-Chancellor, Registrar, Head of Microbiology Department, Dr A Murugan, Supervisor, Periyar University, Salem; DST-FIST (Ref No: SR/FST/LSI-640/2015(C) Dt. 30/5/2016) for the facilities rendered.

---

**Figure 6:**
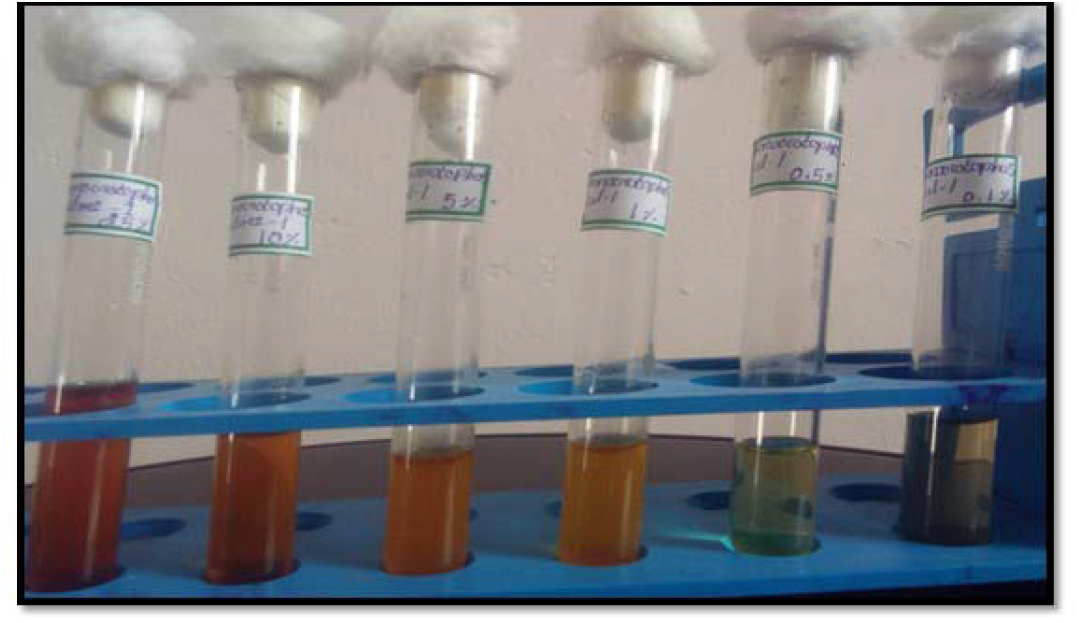
Showing tolerance of isolate V1 to monocrotophos.

**Figure 7:**
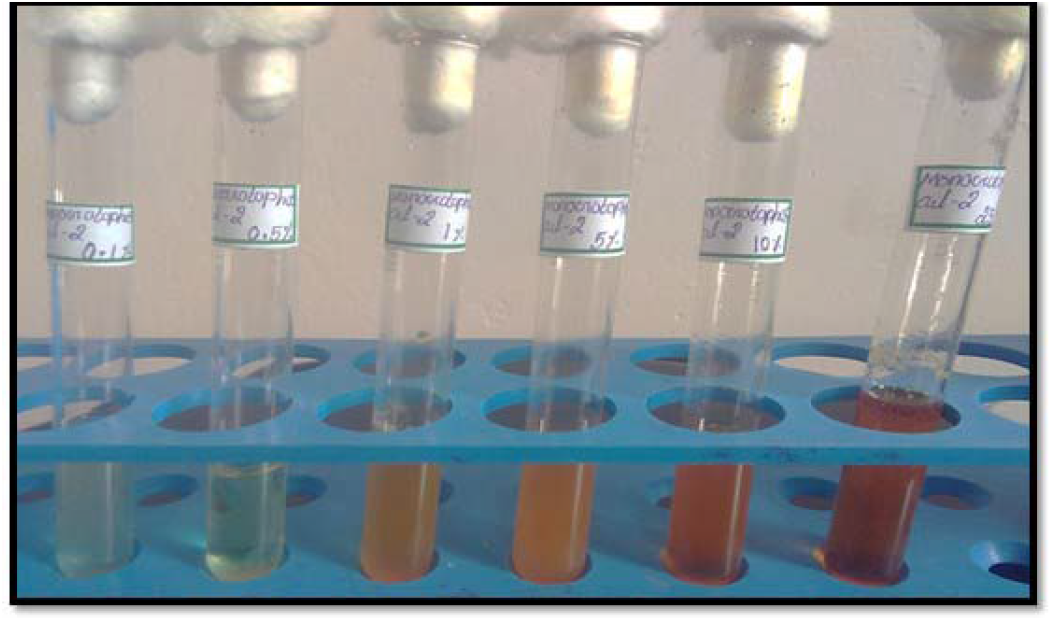
Showing tolerance of isolate V2 to monocrotophos.

**Figure 8:**
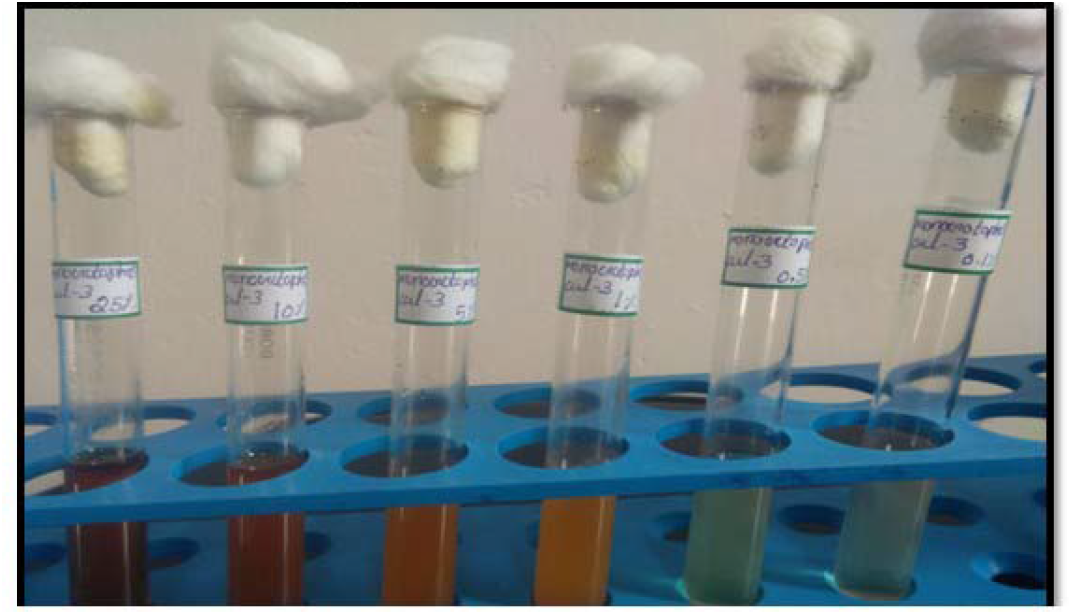
Showing tolerance of isolate V3 to monocrotophos.

**Figure 9:**
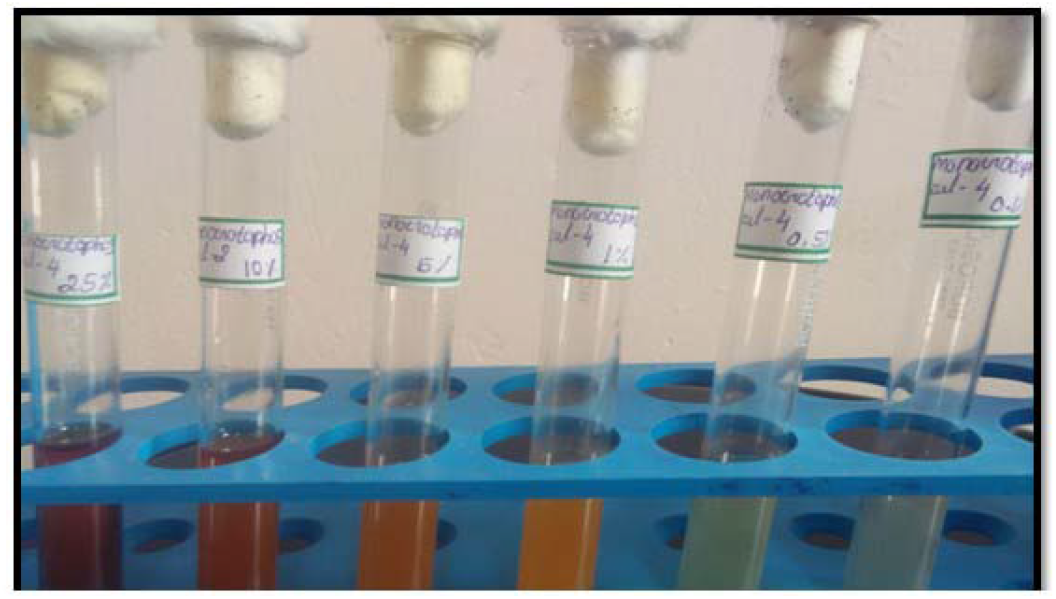
Showing tolerance test of isolate V4 to monocrotophos.

**Figure 10:**
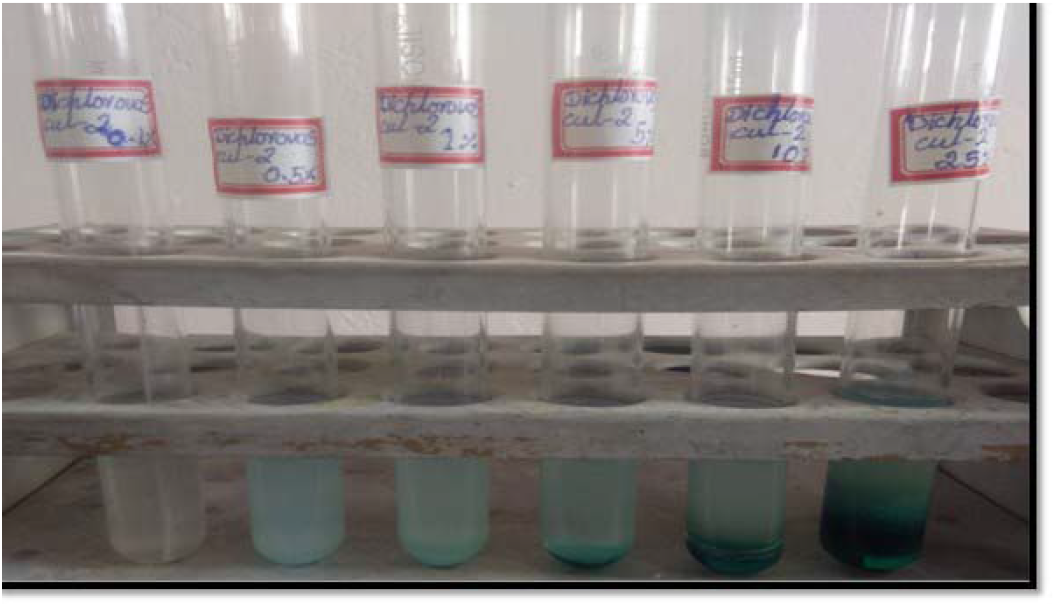
Showing Tolerance Of Isolate V2 To Dichlorovos.

**Figure 11:**
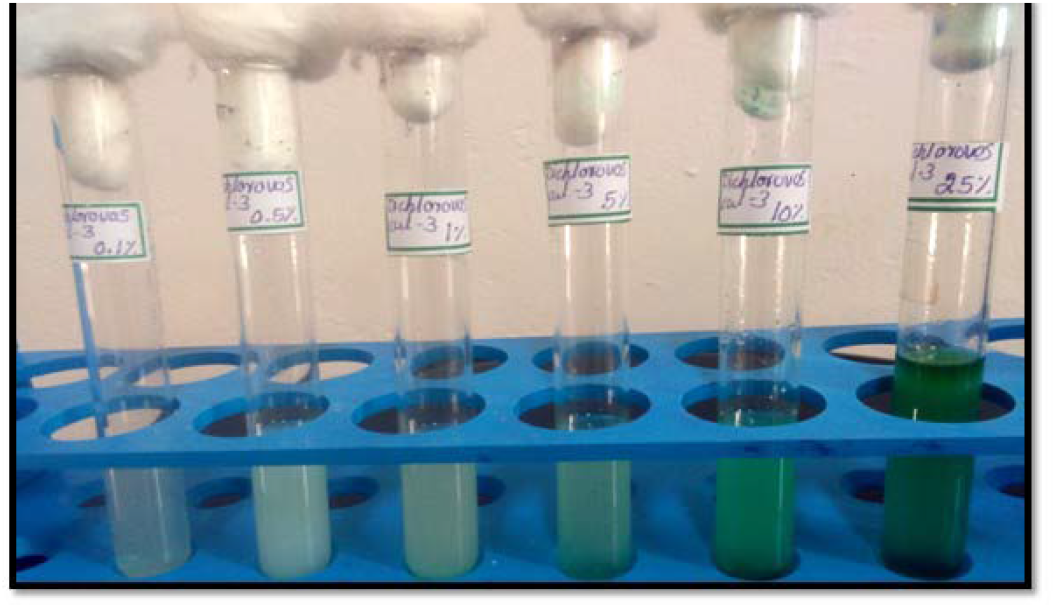
Showing Tolerance Of Isolate V3 To Dichlorovos.

**Figure 12:**
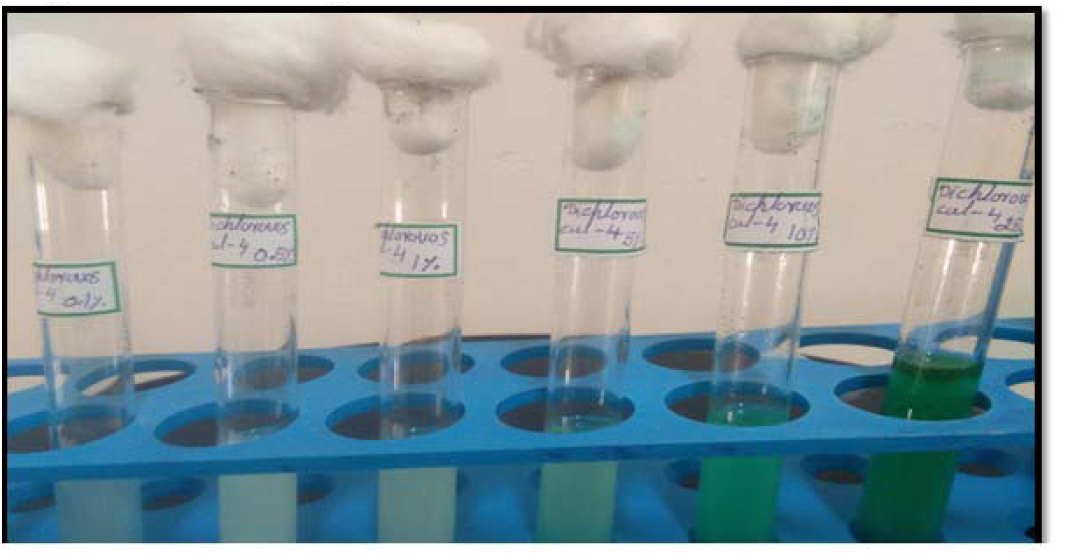
Showing Tolerance Of Isolate V4 To Dichlorovos.

**Figure 13:**
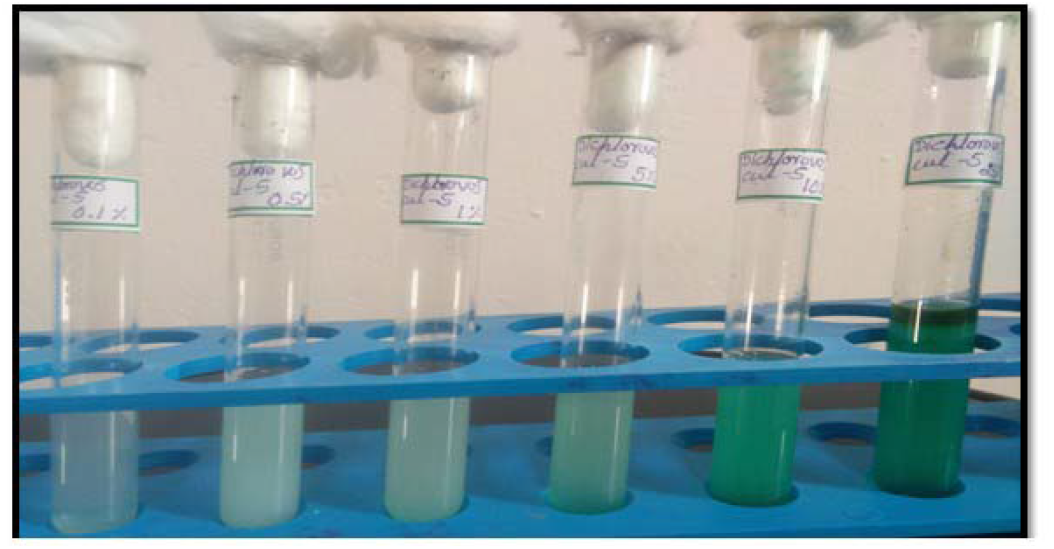
Showing Tolerance Of Isolate V5 To Dichlorovos.

**Figure 15:**
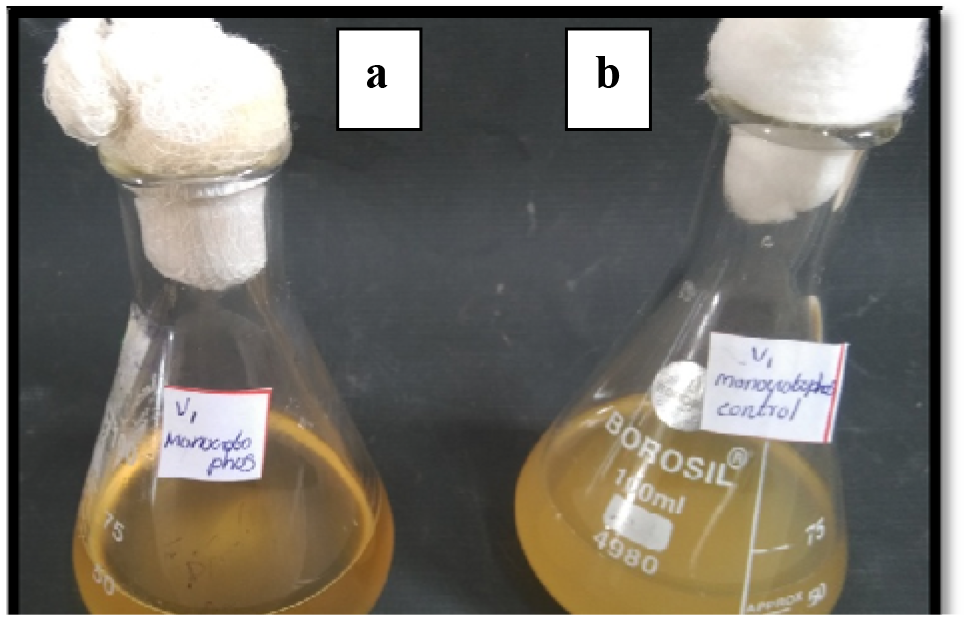
Showing (V_1_) treated with a. 0.1% Monocrotophos and b. control.

**Figure 16:**
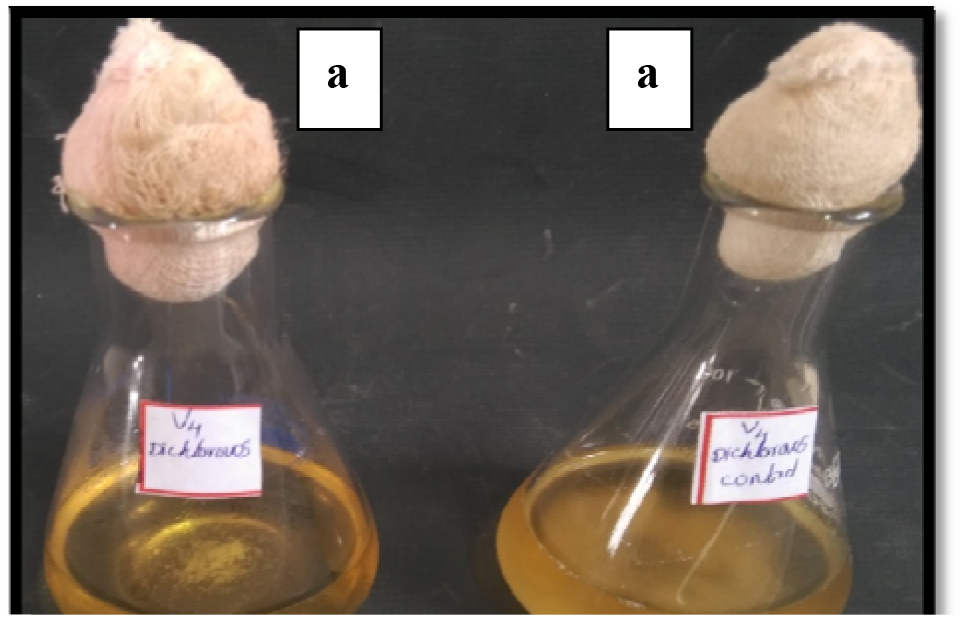
Showing (V_4_) treated with a. 0.1% dichlorovos and b. control.

**Figure 17:**
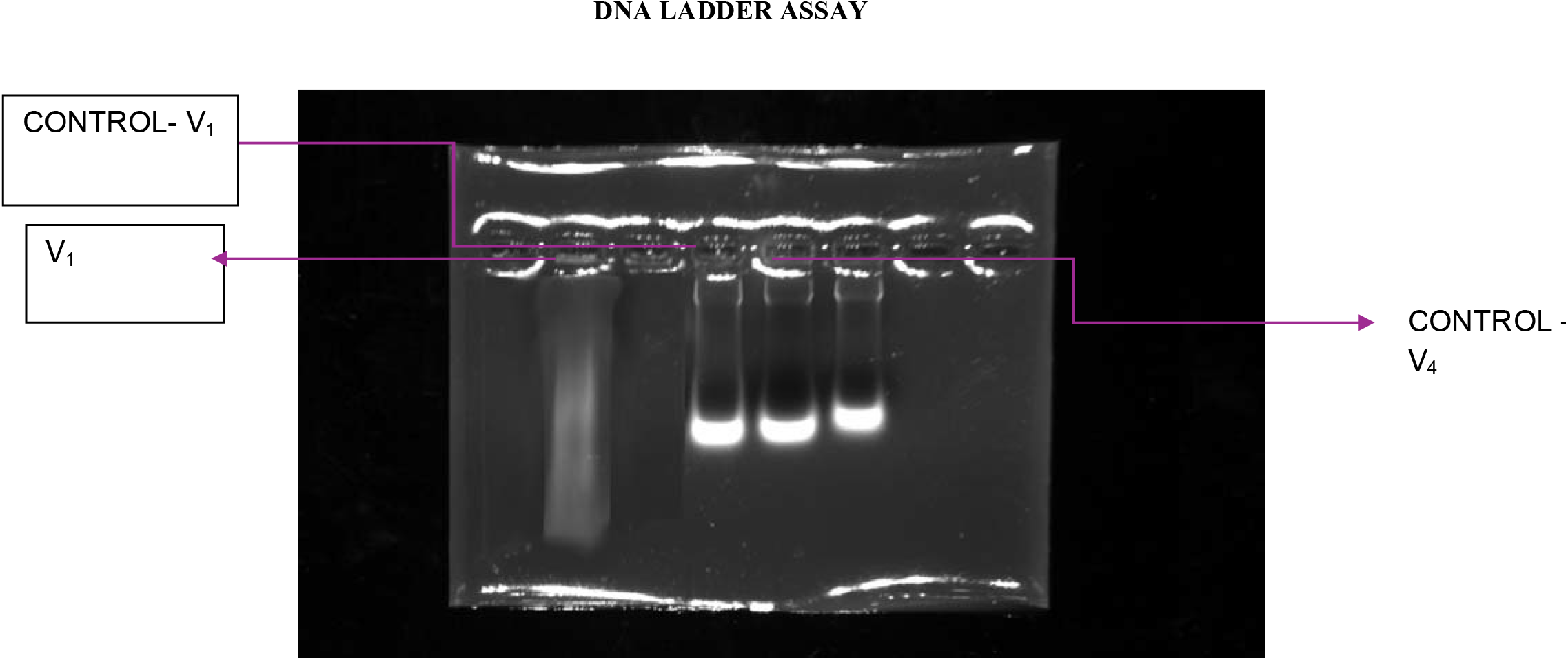
a) Genomic DNA isolation of control culture without pesticide and V1 exposed to pesticide.

**Figure 17 (b).**
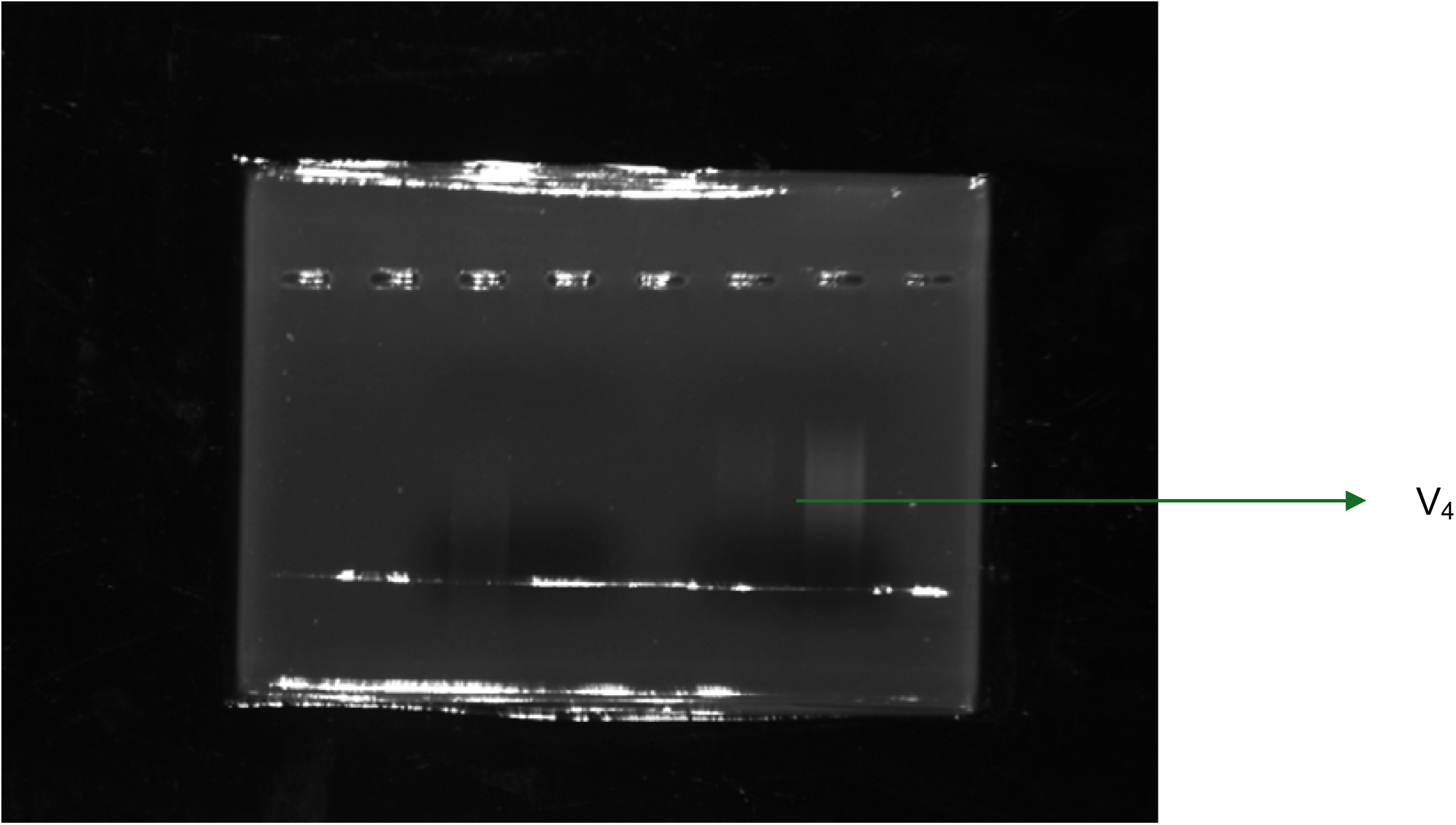
Genomic DNA isolation of culture exposed to pesticides.

